# Integrating Genome-Wide Association and eQTLs Studies Identifies the Genes Associated with Age at Menarche and Age at Natural Menopause

**DOI:** 10.1101/569731

**Authors:** Gang Wang, Jian Lv, Xiaoxin Qiu, Yujun An

## Abstract

**Objective:** An early onset of menarche and, later, menopause are well-established risk factors for the development of breast cancer and endometrial cancer. Although the largest GWASs have identified 389 independent signals for age at menarche (AAM) and 44 regions for age at menopause (ANM), GWAS can only identify the associations between variants and traits. The aim of this study was to identify genes whose expression levels were associated with AAM or ANM due to pleiotropy or causality by integrating GWAS data with genome-wide expression quantitative trait loci (eQTLs) data. We also aimed to identify the pleiotropic genes that influenced two phenotypes.

**Method:** We employed GWAS data of AAM and ANM and Genome-wide eQTL data from whole blood. The summary data-based Mendelian randomization (SMR) method was used to prioritize the associated genes for further study. The colocalization analysis was used to identify the pleiotropic genes.

**Results:** We identified 31 genes whose expression was associated with AAM and 24 genes whose expression was associated with ANM due to pleiotropy or causality. Two pleiotropic genes were identified to be associated with two phenotypes.

**Conclusion:** The results point out the most possible genes which were responsible for the association. Our study prioritizes the associated genes for further functional mechanistic study of AAM and ANM and illustrates the benefit of integrating different omics of data into the study of complex traits.

## Introduction

### Menarche is the first menstrual cycle and signals the possibility of fertility

An early onset of menarche is associated with risks for obesity, type 2 diabetes, cardiovascular disease, breast cancer and all-cause mortality [1]. Menopause is defined as the permanent cessation of menses due to the loss of ovarian follicular activity. Younger age at natural menopause (ANM) is associated with low risk of breast cancer and ovarian cancer, but higher risks of osteoporosis, cardiovascular disease and type 2 diabetes [1]. A Mendelian randomization study have found that later ANM causally increased the risk of breast cancer [2]. These two traits also mark the beginning and the end of a woman’s reproductive life [3].

Genome-wide association studies (GWAS) are capable to identify the association between target phenotypes and million genetic variants. GWAS of age at menarche (AAM) identified 106 loci containing 389 independent signals [4]. GWAS of ANM has successfully identified dozens of loci [2, 5, 6]. Most of these loci encode factors that appear to be involved in DNA repair, immune response and breast cancer processes [2, 5]. However, GWAS can only identify those SNPs strongly associated with target phenotypes, without pinpointing the target genes and the underlying biological mechanism. For example, the largest GWAS of ANM identified 44 loci containing at least one common variant significantly associated with ANM [2]. However, the significant SNPs in 21 loci were annotated to more than one gene in each locus. It suggested that the specific causal genes remain mostly unidentified.

A large part of genetic variants influence the target phenotypes by causal regulatory effect rather than directly influencing the structure of protein [7]. Expression quantitative trait loci (eQTL), which is a genetic variant influencing a target gene’s expression, is often used to explain the underlying biological mechanism of significant SNPs identified by GWAS. Previous studies have suggested that in the significant loci, those SNPs which were also eQTLs were more likely to be functional SNPs [8]. Zhu et al. proposed a summary-based Mendelian randomization (SMR) analysis to combine GWAS and eQTL data into a single analysis [7]. SMR integrates GWAS data and eQTL identified from whole blood tissue to identify potential functionally relevant genes at the significant loci identified in GWAS. Previous studies have shown that whole blood can be a proxy of relevant tissues for various of phenotypes and disease [7, 9].

In this study, we identified genes whose expression levels were associated with AAM or ANM due to pleiotropy or causality, by integrating ANM GWAS data with eQTL data. We conducted a colocalization analysis to identify significant SNPs causally associated with both phenotypes.

## Materials and Methods

### AAM GWAS summary dataset

Using 1000 Genomes Project–imputed genotype data in up to ∼370,000 women, 389 independent signals (P < 5 × 10^−8^) were identified for age at menarche [4]. The summary data were downloaded from the following website (http://www.reprogen.org).

### ANM GWAS summary dataset

The largest-scale GWAS meta-analysis summary data of ANM was used in this study [2]. The GWAS meta-analysis was conducted with a total sample of 69,360 individuals of European descent. SNPs with the minor allele frequency (MAF) no less than 0.01 and the imputation quality larger than 0.4 were included in the meta-analysis. The summary data were downloaded from the following website (http://www.reprogen.org).

### eQTL dataset

Because the Westra eQTL data [9] had a low coverage of human genes, in this study we used the genetic architecture of gene expression (GAGE) eQTL data to do the SMR test [10]. The GAGE study was performed to investigate the genetic architecture of gene expression in peripheral blood in 2,765 European individuals [10]. In the GAGE data, there were 11,829 unique probes. We set the p-value threshold to select the top associated eQTL for the SMR test to be 5×10^-8^. After removing those probes where the p value of the top eQTL was less than 5×10^-8^, there were 8,144 probes left in the eQTL summary data. The binary summary data can be download from http://cnsgenomics.com/software/smr/#DataResource.

### SMR analysis

The method of SMR was fully described in previous paper [7]. In brief, there were three models including causality (Z - > X - > Y), pleiotropy (Z - > X and Z - > Y) and linkage (Z_1_ - > X, Z_2_ - > Y, and Z_1_ and Z_2_ are two variants in linkage disequilibrium (LD) in the cis-eQTL region). In this study, we tried to identify those genes with pleiotropy effect on ANM. To distinguish the causality and pleiotropy model from the linkage model, we conducted the heterogeneity in dependent instruments (HEIDI) test. The null hypothesis of HEIDI test was that there was no heterogeneity in the b_XY_ (the effect of gene’s expression on the target phenotype) values. So we used P_HEIDI_ > 0.05 to exclude those genes belonging to linkage model. The SMR software was downloaded from http://cnsgenomics.com/software/smr/#Download.

### Colocalization analysis

Colocalization analysis was used to identify the genetic variants affecting both phenotypes. The method was detailed in previous paper [11]. In brief, the method based on a hierarchical Bayesian model can be used to find the region containing a variant that influences both phenotypes. It estimates the probability that a given genomic region either 1) contains a genetic variant that influences the first trait, 2) contains a genetic variant that influences the second trait, 3) contains a genetic variant that influences both traits, or 4) contains both a genetic variant that influences the first trait and a separate genetic variant that influences the second trait [11]. The threshold of posterior probability equal to 0.9 was used to control the false discovery rate at level 0.1 [11].

## Results

After removing those probes with the top eQTL at P_eQTL_ smaller than 5×10^-8^, there were 8,144 probes left in the eQTL summary data. The genome-wide significant level for SMR analysis was P_smr_ < 6.14×10^-6^ (0.05/8,144, Bonferroni test). We identified 98 gene-trait associations with P_smr_ < 6.14×10^-6^. After application of the HEIDI test, this reduced to 54 gene-trait associations (P_HEIDI_ > 0.05). Those genes which did not pass the HEIDI test may be associated with AAM or ANM due to linkage.

We identified 31 genes which associated with AAM (Table 1) and 24 genes which associated with ANM due to pleiotropy or causality (Table 2). Three of the 31 genes can be considered as novel, i.e. no previously identified SNP, reported as genome-wide significant in the primary GWAS paper in the cis-eQTL region of the probes. Among the 24 genes, seven genes (MSH6, TLK1, SYCP2L, BRCA1, PGAP3, DIDO1 and DDX17) were previously annotated to be responsible for the association based on distance, biological function, eQTL effect and non-synonymous SNP in high LD. We also identified 5 new genes (AK125462, MSL2, CLSTN3, TRAPPC2L, DDX5 and CPNE1) where there was no significant SNP (p < 5×10^-8^) in the cis-eQTL region of the probes. C17orf46 was the only gene identified to be associated with both phenotypes

**Table 1.**
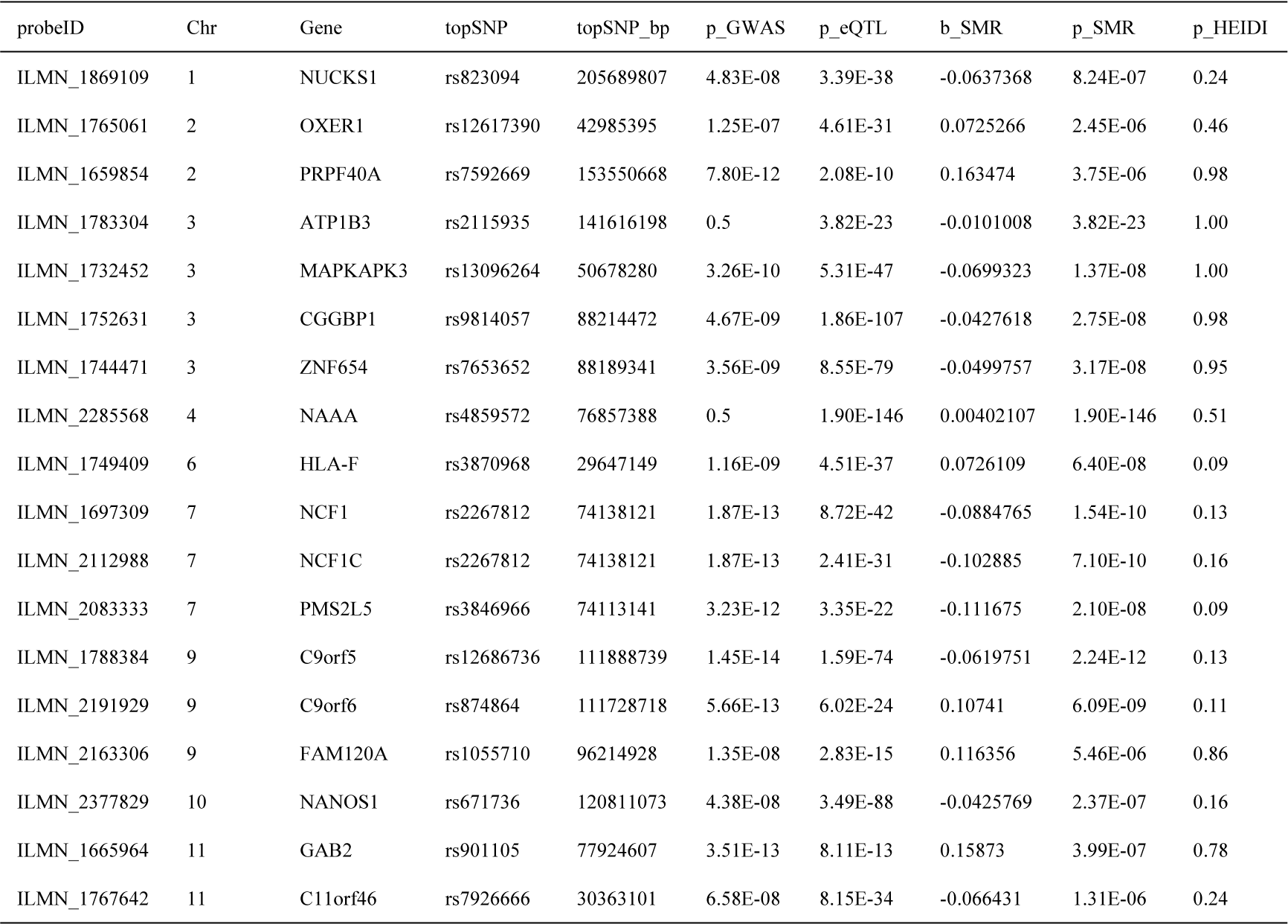
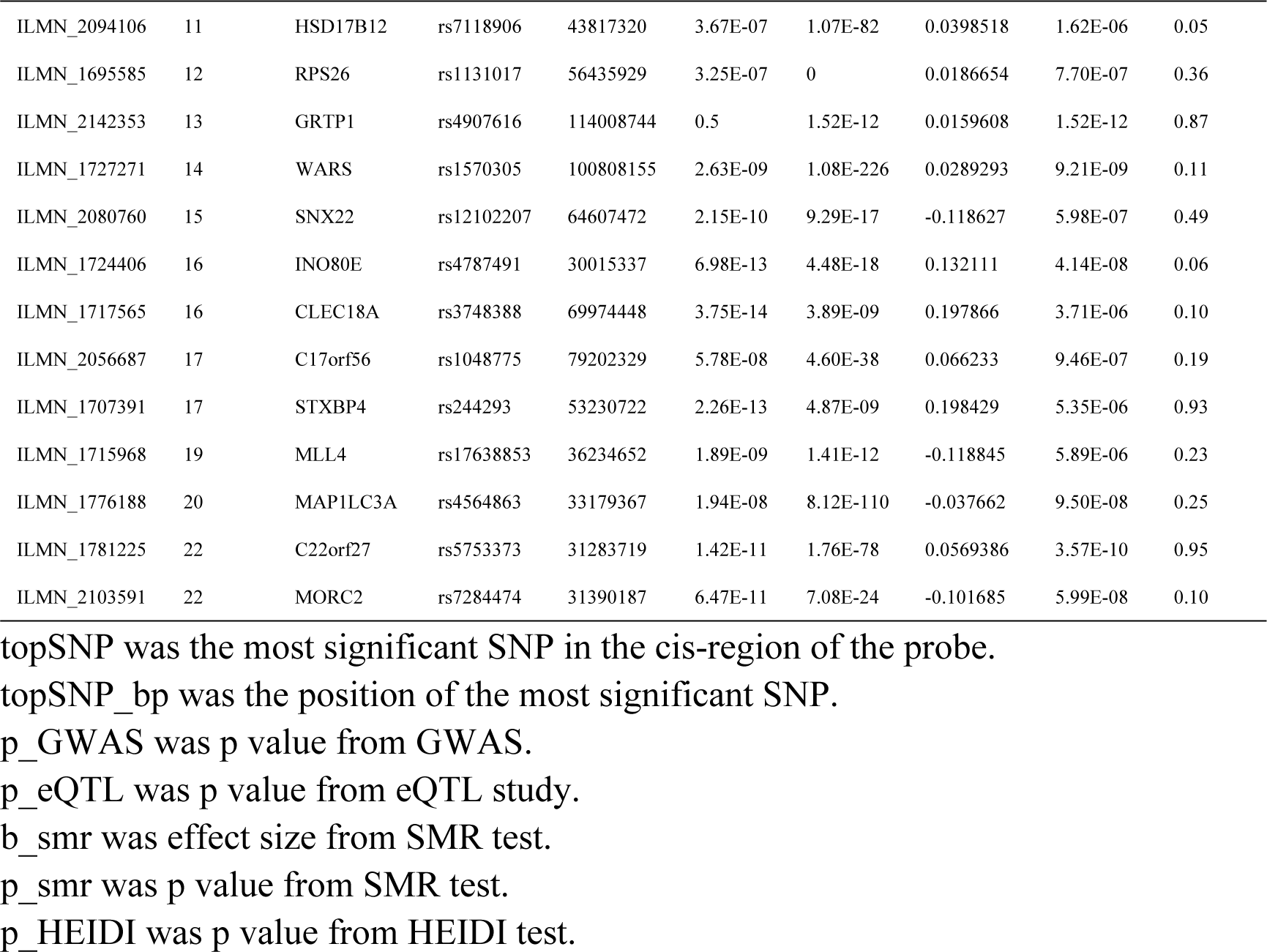
Genes identified by SMR analysis for AAM.

| probeID | Chr | Gene | topSNP | topSNP_bp | p_GWAS | p_eQTL | b_SMR | p_SMR | p_HEIDI |
| --- | --- | --- | --- | --- | --- | --- | --- | --- | --- |
| ILMN_1869109 | 1 | NUCKS1 | rs823094 | 205689807 | 4.83E-08 | 3.39E-38 | -0.0637368 | 8.24E-07 | 0.24 |
| ILMN_1765061 | 2 | OXER1 | rs12617390 | 42985395 | 1.25E-07 | 4.61E-31 | 0.0725266 | 2.45E-06 | 0.46 |
| ILMN_1659854 | 2 | PRPF40A | rs7592669 | 153550668 | 7.80E-12 | 2.08E-10 | 0.163474 | 3.75E-06 | 0.98 |
| ILMN_1783304 | 3 | ATP1B3 | rs2115935 | 141616198 | 0.5 | 3.82E-23 | -0.0101008 | 3.82E-23 | 1.00 |
| ILMN_1732452 | 3 | MAPKAPK3 | rs13096264 | 50678280 | 3.26E-10 | 5.31E-47 | -0.0699323 | 1.37E-08 | 1.00 |
| ILMN_1752631 | 3 | CGGBP1 | rs9814057 | 88214472 | 4.67E-09 | 1.86E-107 | -0.0427618 | 2.75E-08 | 0.98 |
| ILMN_1744471 | 3 | ZNF654 | rs7653652 | 88189341 | 3.56E-09 | 8.55E-79 | -0.0499757 | 3.17E-08 | 0.95 |
| ILMN_2285568 | 4 | NAAA | rs4859572 | 76857388 | 0.5 | 1.90E-146 | 0.00402107 | 1.90E-146 | 0.51 |
| ILMN_1749409 | 6 | HLA-F | rs3870968 | 29647149 | 1.16E-09 | 4.51E-37 | 0.0726109 | 6.40E-08 | 0.09 |
| ILMN_1697309 | 7 | NCF1 | rs2267812 | 74138121 | 1.87E-13 | 8.72E-42 | -0.0884765 | 1.54E-10 | 0.13 |
| ILMN_2112988 | 7 | NCF1C | rs2267812 | 74138121 | 1.87E-13 | 2.41E-31 | -0.102885 | 7.10E-10 | 0.16 |
| ILMN_2083333 | 7 | PMS2L5 | rs3846966 | 74113141 | 3.23E-12 | 3.35E-22 | -0.111675 | 2.10E-08 | 0.09 |
| ILMN_1788384 | 9 | C9orf5 | rs12686736 | 111888739 | 1.45E-14 | 1.59E-74 | -0.0619751 | 2.24E-12 | 0.13 |
| ILMN_2191929 | 9 | C9orf6 | rs874864 | 111728718 | 5.66E-13 | 6.02E-24 | 0.10741 | 6.09E-09 | 0.11 |
| ILMN_2163306 | 9 | FAM120A | rs1055710 | 96214928 | 1.35E-08 | 2.83E-15 | 0.116356 | 5.46E-06 | 0.86 |
| ILMN_2377829 | 10 | NANOS1 | rs671736 | 120811073 | 4.38E-08 | 3.49E-88 | -0.0425769 | 2.37E-07 | 0.16 |
| ILMN_1665964 | 11 | GAB2 | rs901105 | 77924607 | 3.51E-13 | 8.11E-13 | 0.15873 | 3.99E-07 | 0.78 |
| ILMN_1767642 | 11 | C11orf46 | rs7926666 | 30363101 | 6.58E-08 | 8.15E-34 | -0.066431 | 1.31E-06 | 0.24 |
| ILMN_2094106 | 11 | HSD17B12 | rs7118906 | 43817320 | 3.67E-07 | 1.07E-82 | 0.0398518 | 1.62E-06 | 0.05 |
| ILMN_1695585 | 12 | RPS26 | rs1131017 | 56435929 | 3.25E-07 | 0 | 0.0186654 | 7.70E-07 | 0.36 |
| ILMN_2142353 | 13 | GRTP1 | rs4907616 | 114008744 | 0.5 | 1.52E-12 | 0.0159608 | 1.52E-12 | 0.87 |
| ILMN_1727271 | 14 | WARS | rs1570305 | 100808155 | 2.63E-09 | 1.08E-226 | 0.0289293 | 9.21E-09 | 0.11 |
| ILMN_2080760 | 15 | SNX22 | rs12102207 | 64607472 | 2.15E-10 | 9.29E-17 | -0.118627 | 5.98E-07 | 0.49 |
| ILMN_1724406 | 16 | INO80E | rs4787491 | 30015337 | 6.98E-13 | 4.48E-18 | 0.132111 | 4.14E-08 | 0.06 |
| ILMN_1717565 | 16 | CLEC18A | rs3748388 | 69974448 | 3.75E-14 | 3.89E-09 | 0.197866 | 3.71E-06 | 0.10 |
| ILMN_2056687 | 17 | C17orf56 | rs1048775 | 79202329 | 5.78E-08 | 4.60E-38 | 0.066233 | 9.46E-07 | 0.19 |
| ILMN_1707391 | 17 | STXBP4 | rs244293 | 53230722 | 2.26E-13 | 4.87E-09 | 0.198429 | 5.35E-06 | 0.93 |
| ILMN_1715968 | 19 | MLL4 | rs17638853 | 36234652 | 1.89E-09 | 1.41E-12 | -0.118845 | 5.89E-06 | 0.23 |
| ILMN_1776188 | 20 | MAP1LC3A | rs4564863 | 33179367 | 1.94E-08 | 8.12E-110 | -0.037662 | 9.50E-08 | 0.25 |
| ILMN_1781225 | 22 | C22orf27 | rs5753373 | 31283719 | 1.42E-11 | 1.76E-78 | 0.0569386 | 3.57E-10 | 0.95 |
| ILMN_2103591 | 22 | MORC2 | rs7284474 | 31390187 | 6.47E-11 | 7.08E-24 | -0.101685 | 5.99E-08 | 0.10 |

topSNP was the most significant SNP in the cis-region of the probe.
topSNP\_bp was the position of the most significant SNP.
p\_GWAS was p value from GWAS.
p\_eQTL was p value from eQTL study.
b\_smr was effect size from SMR test.
p\_smr was p value from SMR test.
p\_HEIDI was p value from HEIDI test.

**Table 2.**
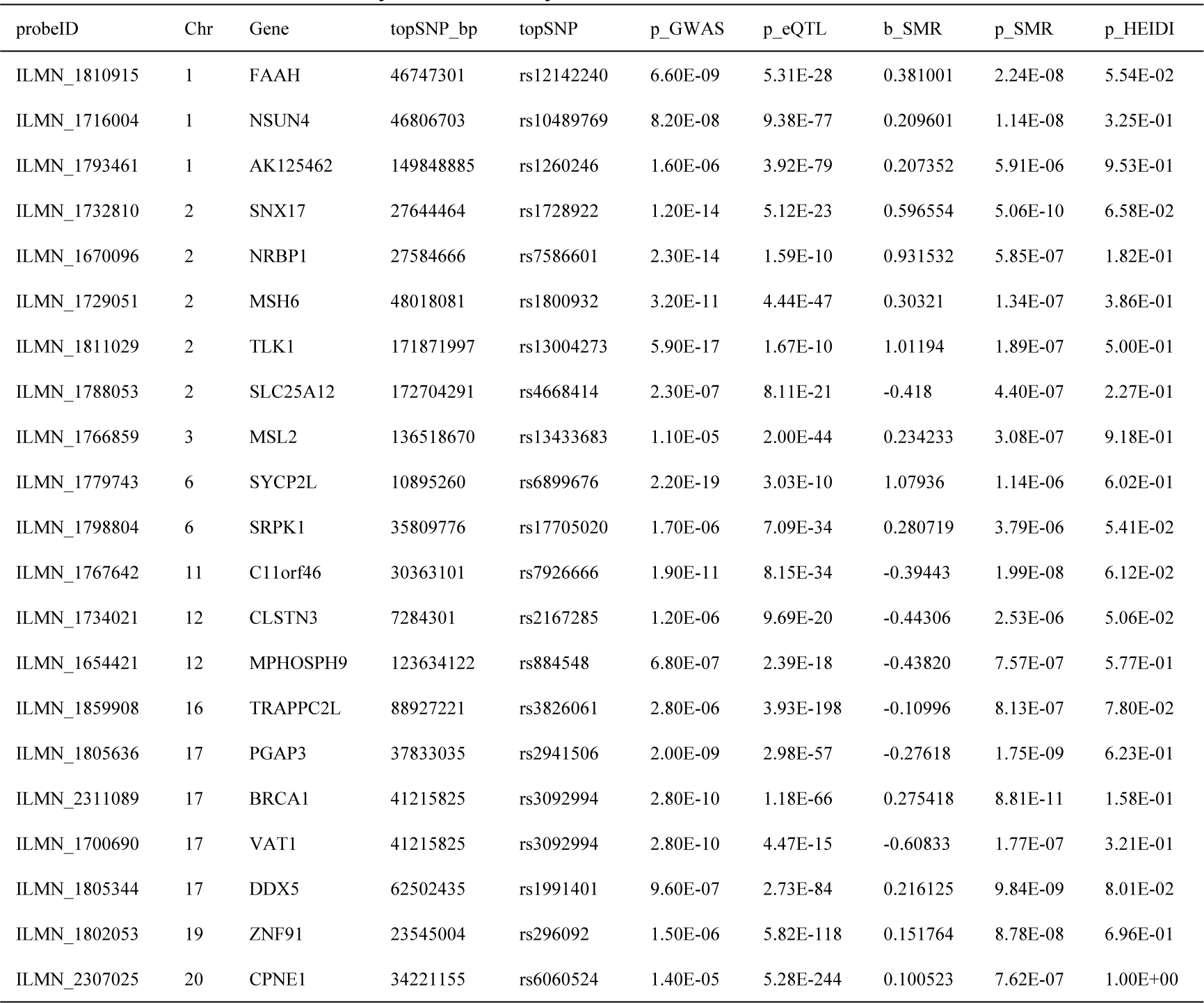
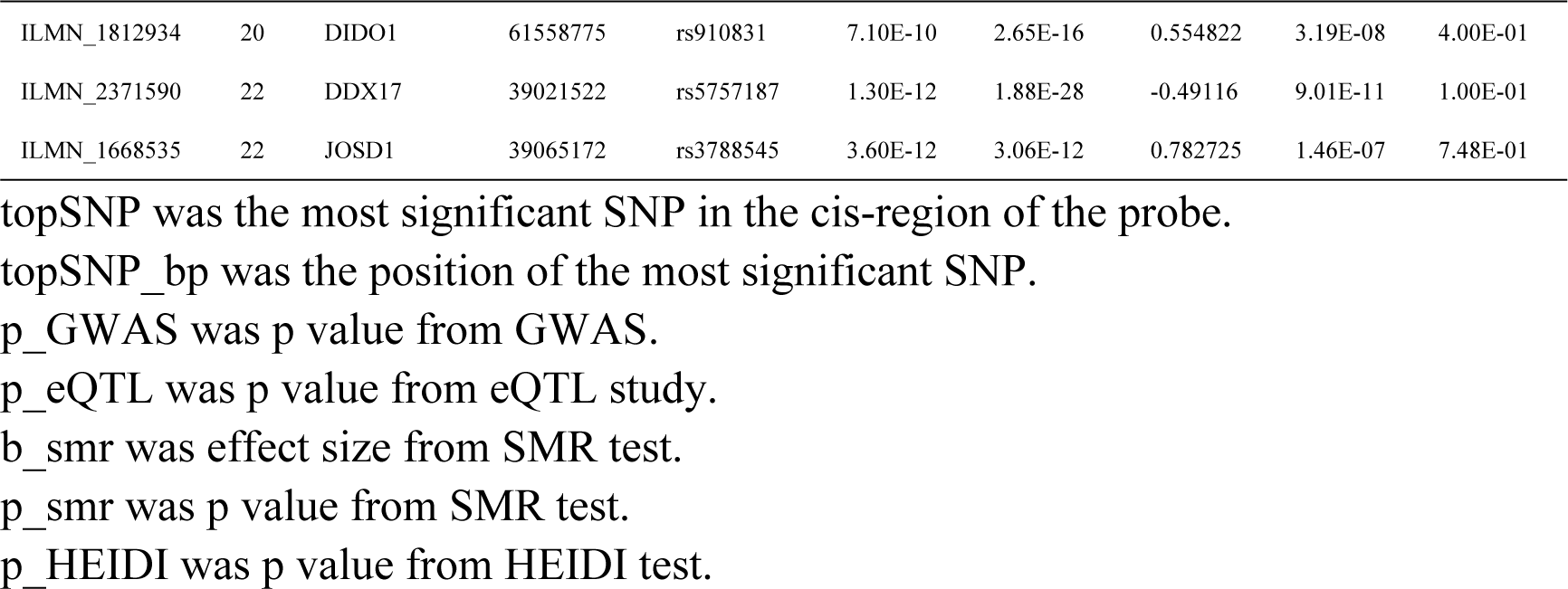
Genes identified by SMR analysis for ANM.

To identify more pleiotropic SNPs and genes associated with both phenotypes, we conducted a colocalization analysis. One region was identified to contain a variant influencing two phenotypes with the posterior probability of 0.92 (Table 3). Thirteen regions were considered to influence two phenotypes through different variants (Table 3). rs3136249, with the largest probability, was considered to be the causal SNP influencing two phenotypes.

**Table 3.** Colocalization analysis results of AAM and ANM.

| chunk | chr | st | sp | PPA_3 | PPA_4 |
| --- | --- | --- | --- | --- | --- |
| 162 | chr2 | 47318990 | 48212562 | 0.92 | 0.064 |
| 1419 | chr15 | 88370262 | 90473690 | 2.5E-17 | 1.00 |
| 1579 | chr19 | 614967 | 2098015 | 1.1E-08 | 1.00 |
| 360 | chr3 | 135458294 | 137370076 | 4.1E-06 | 1.00 |
| 653 | chr6 | 28918936 | 29737846 | 3.3E-05 | 1.00 |
| 799 | chr7 | 92500845 | 93966036 | 5.4E-05 | 1.00 |
| 1637 | chr20 | 32819871 | 34960201 | 6.7E-05 | 0.99 |
| 656 | chr6 | 31572333 | 32682429 | 3.5E-03 | 0.97 |
| 655 | chr6 | 30798697 | 31568469 | 6.1E-04 | 0.94 |
| 1682 | chr22 | 19913726 | 22355640 | 4.0E-04 | 0.94 |
| 128 | chr1 | 241582668 | 242070731 | 2.5E-04 | 0.93 |
| 1644 | chr20 | 42680811 | 44838112 | 1.9E-03 | 0.93 |
| 1693 | chr22 | 37570784 | 39306630 | 3.0E-04 | 0.92 |
| 1213 | chr12 | 55665948 | 57543572 | 3.7E-04 | 0.91 |

Chunk was the internal numerical identifier for the segment.
chr: chromosome
st: star position
sp: end position
PPA\_3 was the posterior probability of model 3.
PPA\_4 was the posterior probability of model 4.

## Discussion

In this study, we identified 31 genes whose expression was associated with AAM and 24 genes whose expression was associated with ANM due to pleiotropy or causality. In total, we identified 7 new genes where there was no significant SNP in the gene region. Many of these genes participated in DNA repair, immune response, and breast cancer process [2, 5]. C17orf46 was identified to be associated with both phenotypes by integrating GWAS and eQTLs data. We also found one region influencing two phenotypes though the colocalization analysis.

SMR demonstrated that it was useful to prioritize novel genes associated with AAM or ANM. In the significant loci, we redefined the functional genes which were more likely to play important roles in the process of natural menopause. For example, a previous study that combined modification quantitative trait locus (mQTL) and eQTL identified that NRBP1 may be a functional gene associated with menopause in the significant locus [12]. Our study found that the expression of NRBP1 was associated with age at natural menopause due to pleiotropy or causality. SRPK1, encoding the splicing factor kinase SRSF protein kinase 1, was highly expressed in basal breast cancer cells [13]. The knockdown of SRPK1 significantly suppressed metastasis of breast cancer cells [14].

SMR tests reduced the multiple hypothesis burdens by testing tens of thousands of genes instead of millions of SNPs [15]. So, SMR was useful in identifying novel genes associated with AAM or ANM. DDX5, which is also known as p68, is a prototypic member of the DEAD box family of RNA helicases that encompasses multiple functions. DDX5 was highly expressed in a high proportion of breast cancers. Patients with a detectable levels of both DDX5 and polo-like kinase-1 (pLK1) often had a poor prognosis [16]. Fatty acid amide hydrolase (FAAH), the enzyme that breaks down the endocannabinoid anandamide and controls its levels, is regulated by estrogen [17].

Despite the common belief that multiple genes are responsible for controlling the timing of menarche and natural menopause, very few genes have been identified that contain common genetic variants associated with AAM and ANM. In this study, we identified two genes rs3136249 is located in the intronic region of MSH6. MSH6, which is a mismatch repair gene, was found to be associated with ANM by previous study [18]. Although the function of C11orf46 is unknown, further studies are needed to prove this result.

The present study may have some limitations that should be considered. Although we redefined the functional genes in the significant loci, these genes may be associated with age at natural menopause due to pleiotropy which meant that some of these genes may be not the causal genes. Due to the limitation of method, we did not distinguish those pleiotropic genes from causal genes. So, further works are warranted to confirm the functional genes and explore the underlying mechanism.

In conclusion, we highlighted the putative functional genes in the significant loci for AAM and ANM. Our study prioritizes the associated genes for further functional mechanistic study of AAM and ANM and illustrates the benefit of integrating different omics of data into the study of complex traits.

## Acknowledgments

We thank Felix R. Day et al. to provide the GWAS summary data of AAM and ANM.

## References

[1] Hartge P. Genetics of reproductive lifespan. Nature Genetics. 2009;41: 637.

[2] Day FR, Ruth KS, Thompson DJ, Lunetta KL, Pervjakova N, et al. Large-scale genomic analyses link reproductive aging to hypothalamic signaling, breast cancer susceptibility and BRCA1-mediated DNA repair. Nat Genet. 2015;47: 1294-1303.

[3] te Velde ER, Pearson PL. The variability of female reproductive ageing. Human Reproduction Update. 2002;8: 141-154.

[4] Day FR, Thompson DJ, Helgason H, Chasman DI, Finucane H, et al. Genomic analyses identify hundreds of variants associated with age at menarche and support a role for puberty timing in cancer risk. Nature Genetics. 2017;49: 834-841.

[5] Stolk L, Perry JR, Chasman DI, He C, Mangino M, et al. Meta-analyses identify 13 loci associated with age at menopause and highlight DNA repair and immune pathways. Nat Genet. 2012;44: 260-8.

[6] Perry JR, Hsu YH, Chasman DI, Johnson AD, Elks C, et al. DNA mismatch repair gene MSH6 implicated in determining age at natural menopause. Hum Mol Genet. 2014;23: 2490-7.

[7] Zhu Z, Zhang F, Hu H, Bakshi A, Robinson MR, et al. Integration of summary data from GWAS and eQTL studies predicts complex trait gene targets. Nat Genet. 2016;48: 481-7.

[8] Schaub MA, Boyle AP, Kundaje A, Batzoglou S, Snyder M. Linking disease associations with regulatory information in the human genome. Genome Res. 2012;22: 1748-59.

[9] Westra HJ, Peters MJ, Esko T, Yaghootkar H, Schurmann C, et al. Systematic identification of trans eQTLs as putative drivers of known disease associations. Nat Genet. 2013;45: 1238-1243.

[10] Lloyd-Jones LR, Holloway A, McRae A, Yang J, Small K, et al. The Genetic Architecture of Gene Expression in Peripheral Blood. Am J Hum Genet. 2017;100: 228-237.

[11] Pickrell JK, Berisa T, Liu JZ, Segurel L, Tung JY, et al. Detection and interpretation of shared genetic influences on 42 human traits. Nat Genet. 2016;48: 709-17.

[12] Zhang X, Moen EL, Liu C, Mu W, Gamazon ER, et al. Linking the genetic architecture of cytosine modifications with human complex traits. Hum Mol Genet. 2014;23: 5893-905.

[13] Gui J-F, Lane WS, Fu X-D. A serine kinase regulates intracellular localization of splicing factors in the cell cycle. Nature. 1994;369: 678.

[14] van Roosmalen W, Le Devedec SE, Golani O, Smid M, Pulyakhina I, et al. Tumor cell migration screen identifies SRPK1 as breast cancer metastasis determinant. J Clin Invest. 2015;125: 1648-64.

[15] Pasaniuc B, L Price A. Dissecting the genetics of complex traits using summary association statistics; 2016.

[16] Iyer RS, Nicol SM, Quinlan PR, Thompson AM, Meek DW, et al. The RNA helicase/transcriptional co-regulator, p68 (DDX5), stimulates expression of oncogenic protein kinase, Polo-like kinase-1 (PLK1), and is associated with elevated PLK1 levels in human breast cancers. Cell Cycle. 2014;13: 1413-23.

[17] Cui N, Wang C, Zhao Z, Zhang J, Xu Y, et al. The Roles of Anandamide, Fatty Acid Amide Hydrolase, and Leukemia Inhibitory Factor on the Endometrium during the Implantation Window. Frontiers in Endocrinology. 2017;8: 268.

[18] Perry JR, Hsu Yh Fau-Chasman DI, Chasman Di Fau-Johnson AD, Johnson Ad Fau-Elks C, Elks C Fau-Albrecht E, et al. DNA mismatch repair gene MSH6 implicated in determining age at natural menopause.

